# Assessment of phylo-functional coherence along the bacterial phylogeny and taxonomy

**DOI:** 10.1101/795914

**Authors:** Marcos Parras-Moltó, Daniel Aguirre de Cárcer

**Affiliations:** Departamento de Biología, Universidad Autónoma de Madrid, Madrid, 28049 Spain

## Abstract

In this report we use available curated phylogenies, taxonomy, and genome annotations to assess the phylogenetic and gene content similarity associated with each different taxa and taxonomic rank. Subsequently, we employ the same data to delimit the frontiers of functional coherence along the bacterial phylogeny. Our results show that within-group phylogenetic and gene content similarity of taxa in the same rank are not homogenous, and that these values show extensive overlap between ranks. Functional coherence along the 16S rRNA gene-based phylogeny was limited to 44 particular nodes presenting large variations in phylogenetic depth. For instance, the deep subtree affiliated to class Actinobacteria presented functional coherence, while the shallower family Enterobacteriaceae-affiliated subtree did not. On the other hand, functional coherence along the genome-based phylogeny delimited deep subtrees affiliated to phyla Actinobacteriota, Deinococcota, Chloroflexota, Firmicutes, and a subtree containing the rest of the bacterial phyla.

**IMPORTANCE:** While bacterial taxonomy and phylogeny resources as well as related bioinformatic tools continue to improve, the question remains as to how they should best be employed in studies using 16S rRNA gene surveys to assess bacteria-ecosystem relationships, a widespread approach. The results contained herein lead to the recommendation that all ranks from genus to class/phylum be employed if using taxonomic binning in the analysis of 16S rRNA gene surveys. With regards to the use of phylogeny or clustering-based approaches, single or arbitrary tree topology or sequence distance thresholds should not be employed. Instead, the results presented here can be used to obtain more meaningful results in many microbial ecology and evolution research scenarios. Moreover, we provide dedicated scripts and files that can be used to continue the exploration of functional coherence along the bacterial phylogeny employing different parameters or input data.

## INTRODUCTION

Our knowledge of microbial communities’ composition has greatly expanded over the last decade thanks to the advent of high-throughput sequencing-based metagenomic approaches. The use of strategies based on the high-throughput sequencing of 16S rRNA gene amplicons is nowadays widespread in studies seeking to assess community structure. The use of such phylogenetic marker is often preferred over shot-gun metagenomic sequencing due to its associated reduced costs and ease of analysis and interpretation. Most commonly, these studies aim not only to provide snapshots of community composition on different samples, but also to assess how these communities change along environmental gradients or sample groups, and significantly, why the observed changes take place in relation to the different putative functional roles of bacterial groups in the ecosystem.

Within these approaches, the taxonomic binning of reads based on the use of dedicated training sets remains the most common strategy in the description of community composition. The use of taxonomic ranks to organize the bacterial tree of life on the basis of evolutionary relationships has a colossal value in scientific communication, and it seems difficult to imagine the progress of microbial ecology without its use. However, the link between taxonomy and phylogeny or ecological function is not clear-cut (1). Another common strategy in the analysis of 16S rRNA gene data is the use of sequence clusters obtained at predefined similarity thresholds (OTUs) or, less frequently yet more meaningfully, the search for study-oriented patterns along the complete 16S rRNA gene phylogeny (e.g. (2, 3)).

The present study analyzes the phylo-functional coherence of taxa and taxonomic ranks, and the frontiers along the bacterial phylogeny where gene content similarity and shared phylogeny cease to correlate. Our main goal is to provide adequate guidance for future analyses employing 16S gene surveys aiming to improve our understanding of the role that Bacteria play in the ecosystem.

## RESULTS

Our results are based on available curated phylogenetic trees and 16S rRNA gene sequence data from the Genome Taxonomy Database (http://gtdb.ecogenomic.org/downloads), as well as genome annotations from proGenomes (http://progenomes.embl.de/). Due to uneven sampling, the number of available members per taxa varied wildly (Suppl. Fig. 1, Suppl. Mat. 1). In all five ranks considered (genus, family, class, order, phylum) a large proportion of groups contained few members, while a small proportion presented a large number of members. This fact will likely impact the overall accuracy of classifiers using the available data as training set (4).

Phylogenetic similarity, assessed using 16S rRNA gene distances as proxy, was also quite variable among groups within the same taxonomic rank (Figure 1A, Suppl. Mat. 1). As expected, the per-rank averages of intra-taxa distance averages followed an ascending trend from genus to phylum (Figure 1A). While correlated, the observed values are markedly smaller than the controversial cut-offs often employed in the field; 0.05, 0.10 and 0.20, for genus, family, and phylum, respectively (5). Another interesting trend relates to the coefficients of variation associated with the per-rank distributions of intra-taxa distance averages (from genus to phylum; 1.08, 0.73, 0.59, 0.48 and 0.36).

**Figure 1.**
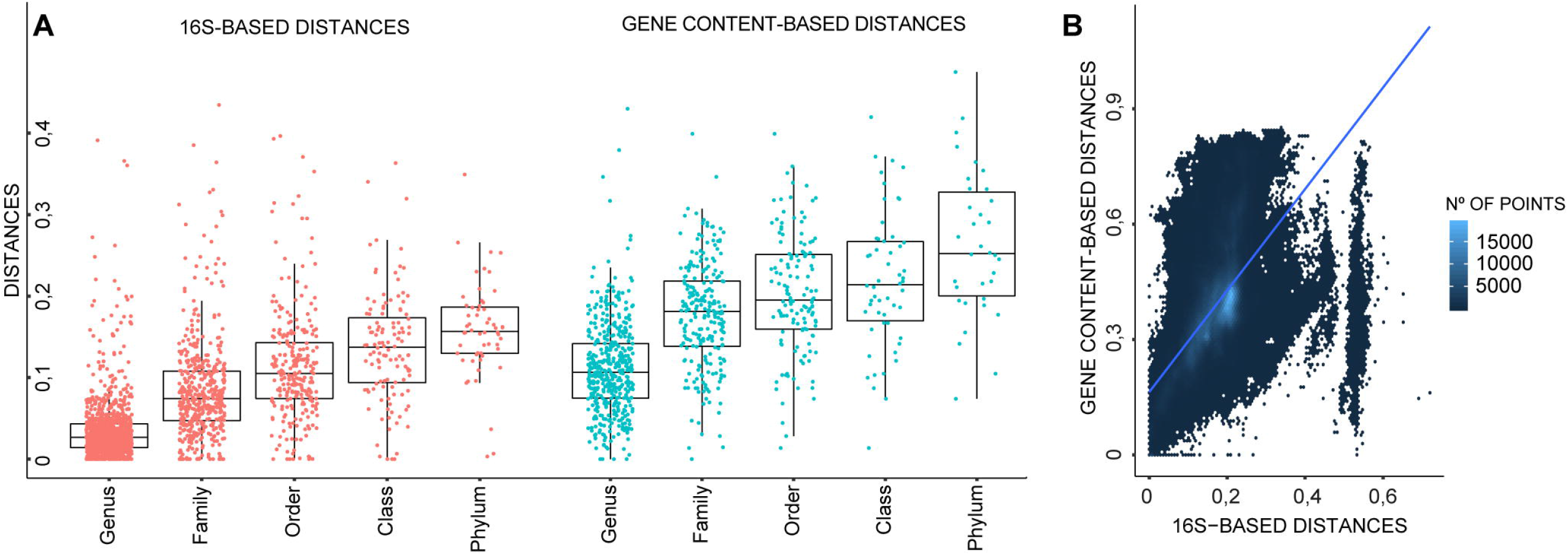
**Panel A;** Box plots describing average within-group phylogenetic and gene-content distance values for each of the proposed taxa in all considered ranks. **Panel B;** Correlation between 16S rRNA gene and gene content-based distances. Each point represents both distance values for each of the possible pairwise comparisons among the 6 989 dereplicated genomes with genome annotations (y = 1.32x + 0.16, R^2^ = 0.37).

The values seem to indicate that phylogenetic depth of taxa in a rank becomes more stable from genus to phylum. Equally anticipated, per-rank distributions of intra-taxa gene content distance averages also followed an ascending trend from genus to phylum (Figure 1A, Suppl. Mat. 1).

When analyzing intra-taxa distance averages (Figure 1A) and standard deviations (Suppl. Fig. 2) it becomes clearer that taxonomic ranks are not cohesive in terms of phylogenetic depth and gene content similarity. That is, for all taxonomic ranks, the different taxa within the same rank do not present a homogeneous level of either phylogenetic depth or gene content similarity. In this regard, the phylogenetic and gene content similarity values of many taxa was lower than those for taxa in superior ranks, and *vice versa*. This circumstance most likely stems from the fact that available genomes are not evenly distributed throughout the bacterial tree of life, that for operational and communication purposes the number of ranks is small, and that the main focus of modern taxonomy is related to the production of monophyletic taxa (6).

Our initial results showed the expected overall correlation between gene content and 16S-based distances (Figure 1B). However, a noticeable discontinuity appeared in the pairwise 16S-based distance values at *ca*. 0.5 which, in turn, seems to mark the limit of the observed correlation. In order to better explore the frontiers of functional coherence along the bacterial phylogeny, we developed *FunCongr.R;* a script that estimates functional coherence at each node of a tree by comparing within-node average gene content distance against what could be expected from a random draw of genomes along the phylogeny. The script also defines a node as non-functionally-coherent if any of its descendant nodes is non-functionally-coherent.

The analysis of the 16S rRNA gene phylogeny (Figure 2, Suppl. Fig. 3, Suppl. Mat. 2) returned 44 functionally-coherent nodes. These nodes present large variations in depth, showing within-node average 16S rRNA gene distance values ranging from 0.03 to 0.25 (average 0.12±0.05). Without the goal of being exhaustive, a subsequent analysis based on the consensus (80%) taxonomy of leaves in each node, with the sole criterion of whether a particular taxon appears assigned to a single coherent node, revealed various deep-branching nodes affiliated to the Acidobacteriota (p), Alphaproteobacteria (c), Actinobacteria (c), Bacilli (c), Bacteroidia (c), Cyanobateriia (c), Fusobacteriales (o) Campylobacterales (o), and a node (node2739) dominated by sequences affiliated to Bacilli (c) and Clostridia (c) (61% and 22%, respectively). On the other hand, the Gammaproteobacteria-affiliated region of the tree was partitioned among many different coherent nodes, hence indicating that such class is not functionally coherent in itself. Here, the Betaproteobacterales (o) stood out as a large and coherent node, while many different coherent nodes appeared affiliated to the Pseudomonadales (o), Enterobacterales (o), and even Enterobacteriaceae (f). On the other hand, the analysis of the genome-based phylogeny (Suppl. Fig. 4-5, Suppl. Mat. 2), inferred from a concatenated alignment of 120 ubiquitous single-copy proteins (6), returned four functionally-coherent deep subtrees, respectively affiliated to phyla Actinobacteriota, Deinococcota, Chloroflexota, Firmicutes (node15643), and a fifth functionally-coherent deep subtree containing the rest of the bacterial phyla (node6).

**Figure 2.**
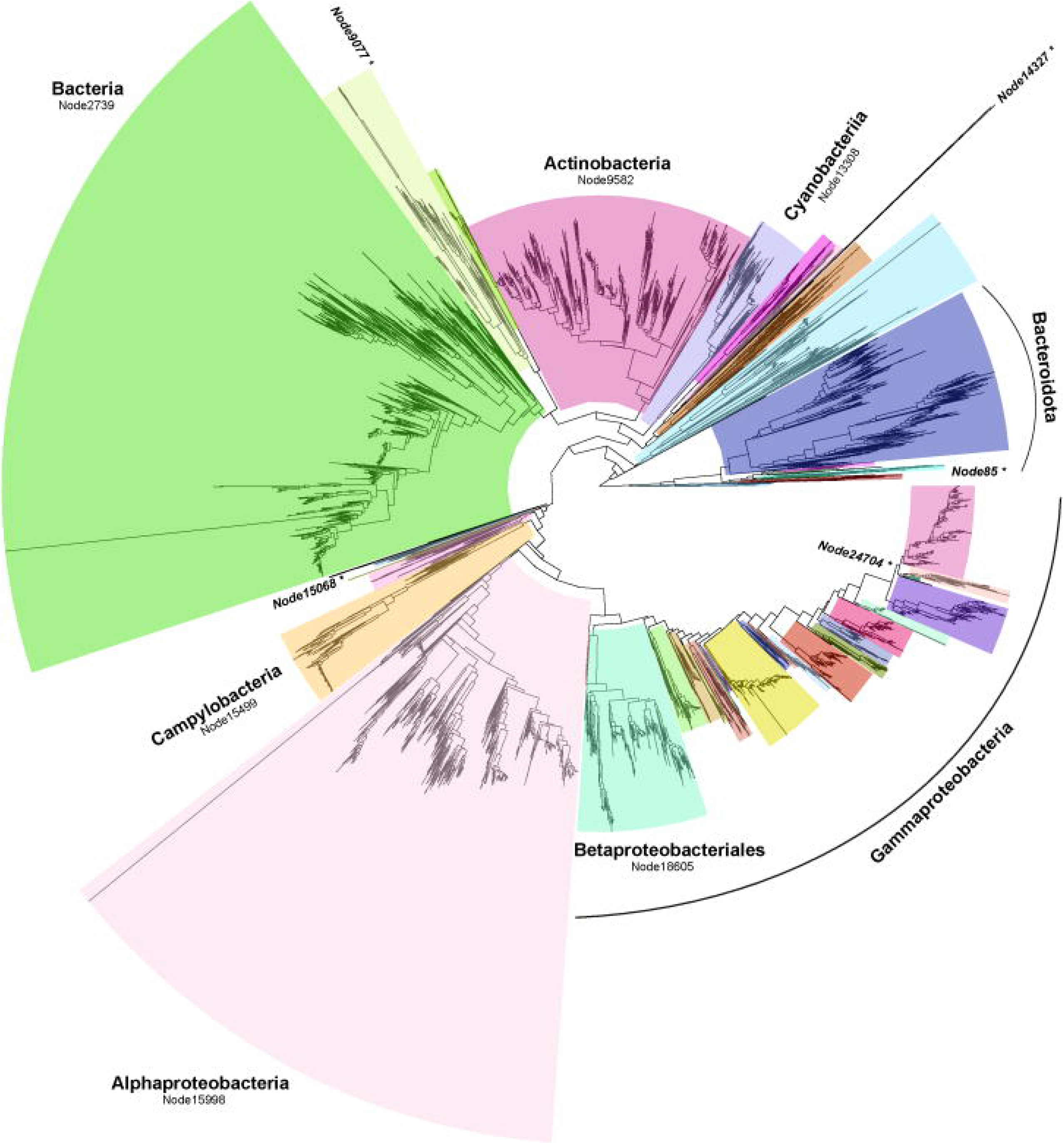
Frontiers of functional coherence along the 16S rRNA gene phylogeny. The tree represents Park *et al*’s 16S rRNA gene-based phylogeny, pruned to contain only leaves with nearly full length 16S rRNA sequences and functional annotations. Colored boxes mark limits of phylo-functional coherence. Only the most prominent results are annotated (see Suppl. Fig. 3 for an annotated and vertical version of the tree). *Nodes that failed to past the functional coherence test.

## DISCUSSION

While bacterial taxonomy and phylogeny resources, as well as related bioinformatic tools, continue to improve, the question remains as to how they should best be employed in studies using 16S rRNA gene surveys to assess bacteria-ecosystem relationships, a widespread approach. With regards to the use of taxonomic binning, our results show that within-group phylogenetic and gene content similarity of taxa in the same rank are not homogenous, and that these values show extensive overlap between ranks. Thus, we believe that most taxonomic ranks should be considered when assessing bacteria-ecosystem relationships, since it is not possible to know *a priori* which sections and depths of the bacterial phylogeny will be affected by the studied environmental gradient or sample group. The possible exceptions are species and phylum, since the 16S rRNA gene presents low phylogenetic resolution at the highest and lowest taxonomic ranks (7), and very few of the dereplicated genomes employed by Parks *et al*. present species-level classification.

With regards to the use of phylogeny-based approaches, the 16S rRNA gene-based phylogeny showed remarkable correlation with gene content similarity. However, the breaking points of such relationship appeared at different depths along each different branching path of the phylogeny. Thus, single, arbitrary tree topology or sequence distance thresholds should not be employed regarding phylo-functional coherence. For instance, while a deep Actinobacteria (c)-affiliated node passed our functional coherence test, the Enterobacteriaceae (f)-affiliated shallower subtree did not.

The strategy employed could be biased by low quality genome annotations, which prompted us to employ the curated proGenomes database. In this sense, the depth of the frontiers of functional coherence described here could be understood as conservative estimates; for instance within the Bacteroidota (p)-affiliated subtree of the 16S-based phylogeny, node85 failed to pass the functional coherence test, and thus the subtree was split into several coherent units (Figure 2, Suppl. Fig. 3). While reducing the p-value of the permutation test to 0.01 returned the exact same tree topology, node85 contains only five genomes, and thus missannotation could potentially have artificially reduced the limits of phylo-functional coherence in this subtree. On the other hand, within that same phylogeny the loss of functional coherence within the Enterobacteraceae (f)-affiliated subtree, and hence that of the larger Gammaproteobacteria (c)-affiliated subtree as well (Figure 2, Suppl. Fig. 3), was supported by the analysis of 35 genome annotations (node24704).

In addition to our previous results-driven recommendation that all ranks from genus to class/phylum be employed if using taxonomic binning, we argue that our functional coherence proxy (i.e. more within-node gene content similarity than expected by chance, and no descendant nodes failing to pass such test) is intuitive and useful. Thus, the results presented here can be used by the community, together with Park *et al*.’s provided phylogenetic trees, to obtain more meaningful results in many microbial ecology and evolution research scenarios. Moreover, the associated scripts and files are freely available (https://git.io/Jec5U) so that (e.g.) different phylogenies, genome annotations, or significance thresholds can be used in the assessment of phylo-functional coherence.

## METHODS

### Phylo-functional coherence of taxa and ranks

First, the 16S rRNA gene sequences for the representative (dereplicated) genomes employed to produce Parks *et al.*’s taxonomy (bac_ssu_r86.1.fna) were obtained from their repository (http://gtdb.ecogenomic.org/downloads)(6). Then, *Mothur* (8) commands were used to align the sequences against SILVA reference alignment (9), and trimmed to contain only the sequence delimited by universal primers 27f and 1492R (10). Sequences not spanning the region were removed, leaving a total of 15 186 sequences. Taxa not including at least three members were not considered for subsequent analyses. Distances between the remaining 16S rRNA gene sequences were obtained using *Mothur*, and later analyzed on the basis of their taxonomic affiliation (bac_metadata_r86.tsv), producing summary statistics for all taxa at all ranks.

NCBI’s TaxIDs for each of the representative genomes (included within bac_metadata_r86.tsv) were linked to the gene content information obtained from the proGenomes resource (11) (complete annotation table kindly provided by Daniel Mende), resulting in 6 989 different annotations for representative genomes presenting nearly full length 16S rRNA gene sequences. Pairwise Jaccard distances between these genomes were derived from the shared presence of genes using the *R* package *vegan*, again producing within-taxa summary statistics for all taxa at all ranks.

### Frontiers of functional coherence within phylogenetic trees

We developed a dedicated R script, *FunCongr.R*, to explore the limits of functional coherence along the bacterial phylogeny. Our approach employs phylogenetic trees (gtb_r86.ssu.bacteria.fasttree.tree and bac120_r86.1.tree (6)) and pairwise Jaccard distances between genomes obtained as mentioned above. The trees were initially processed to include constant node labels, and pruned to remove leaves without functional annotation.

The script traverses the tree from leaves to root in such a way that nodes are evaluated only if its descendant nodes have already been evaluated. Only nodes with at least 5 leaves are analyzed. At each non-flagged node, the script compares the average value of pairwise Jaccard distances between its leaves to an empirical cumulative distribution (ECD) of 1000 average values of pairwise Jaccard distances between N random leaves along the phylogeny, where N is the actual number of leaves of the node being evaluated. If the node’s average Jaccard distance value is not within the lowest 5% of the ECD, the node is flagged. Once a node has been flagged, all its ascendant nodes are automatically flagged.

In this manner, we delimit functional coherence along the tree by pinpointing nodes that either i) fail to pass the random ECD test, or ii) present descendant nodes that fail to pass the ECD test. Thus, here functional coherence is presumed to persist if within-node average gene content distance is lower (i.e. more similar) than what could be expected from a random draw of genomes along the phylogeny (with p<0.05). The results are finally processed by recording the last functionally-coherent node along each branching path, along with their summary statistics. All figures of this work were produced using *ggplot2* package in R.

## Acknowledgements

We thank the Bioinformatics Unit at CBMSO for their support. This work was funded by the Spanish Ministry of Science and Innovation grant BIO2016-80101-R. The funders had no role in study design, data collection and interpretation, or the decision to submit the work for publication

## Data availability

The datasets analyzed during the current study are available from their original source (as stated above) and at (https://git.io/Jec5U).

