## Supplementary figures 1-2 for "Assessment of phylo-functional coherence along the bacterial phylogeny and taxonomy"

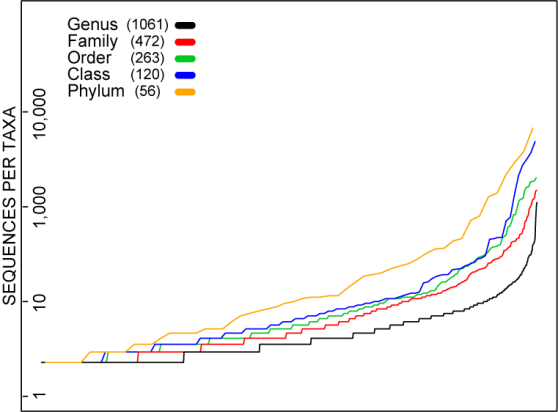

**Supplementary Figure 1.** Rank abundance curves describing the number of members for each of the proposed taxa in all considered ranks.

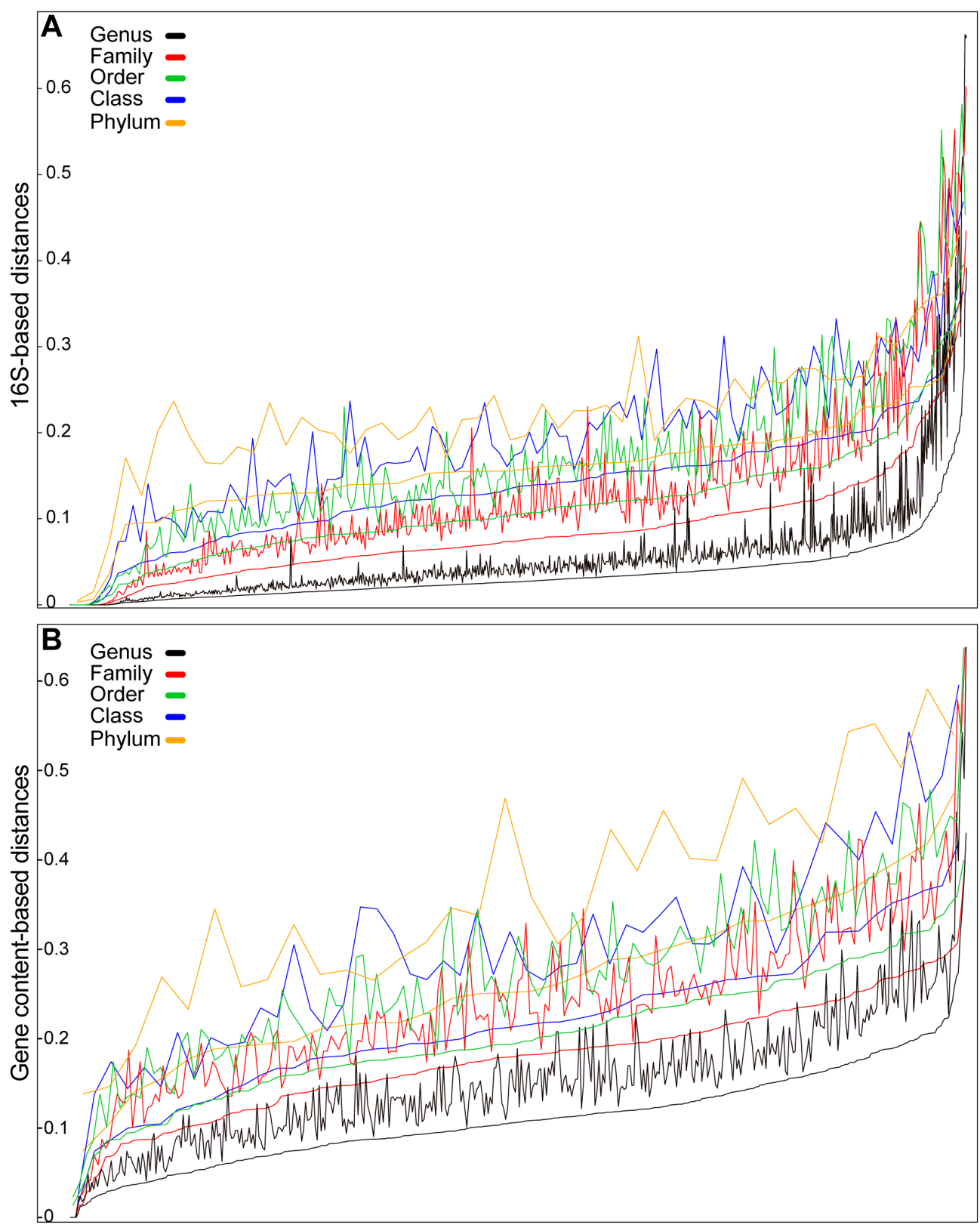

**Supplementary Figure 2.** Rank curves describing the within-group average and average plus standard deviation phylogenetic (A) and gene-content (B) distance values for each of the proposed taxa in all considered ranks.
