## Supplementary figures and images for "Assessment of phylo-functional coherence along the bacterial phylogeny and taxonomy"

### Supplementary figure 3

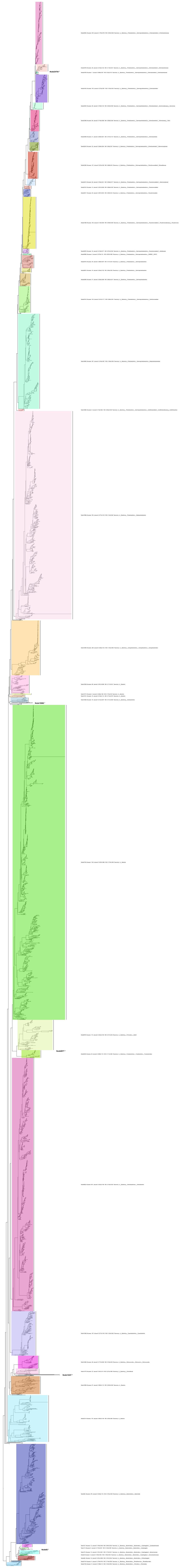

### Supplementary figure 4

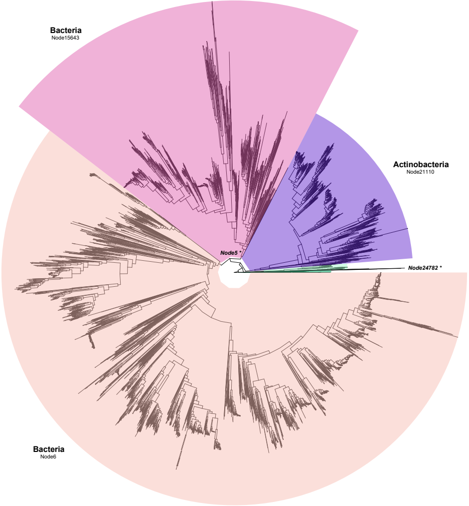

### Supplementary figure 5

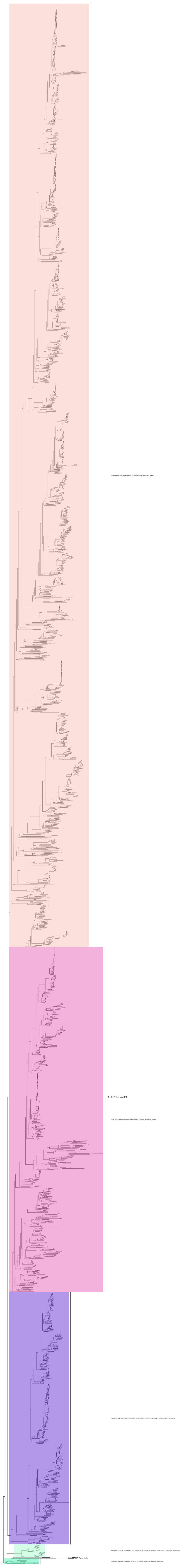
